# GO, NO-GO: A SENSITIVE PERIOD FOR DEVELOPING SEX-SPECIFIC VISUALLY-GUIDED PREY PURSUIT-PREDATOR AVOIDANCE TRADE-OFF STRATEGIES

**DOI:** 10.64898/2026.08.14.744940

**Authors:** Arnab Biswas, Ting Feng, Rocio Olvera, Geo January, Jennifer L. Hoy

## Abstract

In natural environments, animals face a trade-off between foraging and staying vigilant against predators—a conflict known to shape sensory processing and behavior through natural selection. Even within a species, this trade-off may be resolved differently depending on age, sex, or life history. To identify the neural mechanisms underlying visually-guided pursuit-avoidance trade-offs, we studied how mice responded to a sudden overhead threat while pursuing a moving, prey-like target near the ground. This paradigm let us quantify orienting decisions in adolescent and adult mice of both sexes. We found that adolescence is a key period when male and female mice begin to diverge in how they resolve this conflict. Manipulating the value of the pursued target during adolescence versus adulthood revealed that adolescence is also a sensitive period for shaping adult escape-to-shelter versus continue to approach target behavior. Notably, hunting experience gained specifically during adolescence caused males—but not females—to shift strategy: experienced adolescent males tended to uniquely shift towards a "no-go" (stay-near-Prey) strategy. All the other groups instead favored an active "go" strategy, repeatedly shifting between approaching target and running to shelter. All mice showed this oscillation between approaching target and escaping to a shelter to some degree, but hunting experience during adolescence most significantly shifted this balance between approach and escape for males.

## Introduction

Studying visual loom-evoked responses in mice has revealed key neural mechanisms controlling threat orienting. Innate looming-evoked orienting is conserved from insects (Card, 2012) to primates (Schiff et al., 1962) (see Peek & Card, 2016; Branco & Redgrave, 2020, for reviews), and is plastic and life-history dependent in mammals (Albrecht et al., 2025).

Substantial progress has been made in understanding the neural circuits underlying looming-evoked avoidance in adult mice (Evans et al., 2018; Shang et al., 2018; Li et al., 2023). However, few studies address how these behaviors develop in more natural contexts that reflect a species’ sensory ecology. For example, mice and other species would rarely forage far from the safety of a shelter nor explore open environments in the absence of a motivating reward (Lai et al., 2024; Thompson et al., 1982). Similarly, all species have evolved trade-offs between responding to visual threats and exploiting visually-targeted resources (Lima & Dill, 1990). Indeed, head-position and posture observations of animals foraging under threatening and non-threatening conditions show that vigilance is reduced and predator detection is impaired when animals lower their heads to feed, pursue their own prey or engage with conspecifics (Lima & Dill, 1990; Linson et al., 2007). These foraging-vigilance trade-offs are known to depend on sex (Garcia M. et al., 2023). Yet the neural wiring differences that give rise to such life-history-dependent variation remain unidentified— an important gap in understanding how animals achieve flexible, efficient orienting and avoid debilitating anxiety and fear.

To address this gap, we developed a novel "approach-avoidance conflict" paradigm in which competing-valence visual stimuli co-occur as they would in nature. This lets us study the development of visually guided choice behavior in an ecologically relevant, evolutionarily shaped context, and in turn understand conserved strategies animals use to balance resource exploitation against predation risk. In this paradigm, we first capture mice’s attention and positive orienting responses using sweeping motion stimuli near the arena floor. Once the animal begins approaching, we trigger an overhead loom mimicking a predator’s approach—creating a simultaneous conflict between threat and reward. Notably, ambush predators from snakes to cats exploit this same conflict in their visually sensitive prey (birds and mice) through caudal luring or tail-flicking, a form of aggressive mimicry that increases hunting success (Bostanchi et al., 2006; Hagman et al., 2008). Thus, it is highly likely that our observations relate directly to those predators most salient to mouse/rodent ecology.

In contrast, other studies of loom-evoked evasion in mice typically present the overhead loom while animals freely navigate an open environment (Yilmaz et al., 2013), consume spatially localized food or water (Kast et al., 2026), or process overhead sweeping stimuli as a separate threat cue (Baier et al., 2025). These contexts are well-characterized mechanistically, but evasive choice within them varies considerably with lab environment, prior experience, sex, age, and evolutionary history (Baier et al., 2025; De Franceschi et al., 2016; Salay et al., 2018, Shang & Liu et al., 2015), producing variable rates of freezing versus escape-to-shelter and avoidance behavior dynamics. Most studies that trigger the loom when a mouse spontaneously enters a given region assume the stimulus engages the upper visual field via the lower retina—but this is rarely verified, and eye-head coupling during exploration is in fact complex and variable for mice (Benquet et al., 2026). Further, observations of rodents and other species in natural environments suggests that head position is key to indicating the relative state of vigilance and responsiveness to predator cues of foraging animals (Makowska and Kramer, 2007; Linson et al., 2026; Wallace et al., 2013). This feature of animal behavior, head or eye position, has not been used systematically in prior studies to align responses to predatory cues. Mice hunting crickets, for instance, lower and angle their heads to fixate ground-level prey within the middle-to-lower retina, maximizing binocular overlap near the target (Johnson et al., 2021). Other work shows binocular vision facilitates escape via superior colliculus circuits in the same mouse strain, C57BL/6 (Broersen et al., 2025). Together, this suggests overhead looming or sweeping stimuli may compete with ground-level stimuli for binocular visuospatial resources. We therefore modified our existing computerized spontaneous perception of objects task (C-SPOT; Procacci et al., 2020) to deliver overhead looming threats only once mice were already accurately pursuing ground-level, prey-like sweeping stimuli—ensuring the loom reliably appears in the upper/peripheral visual field while the mouse is engaged in binocular processing of a ground-level target.

Using this modified C-SPOT paradigm, we found that mice rarely escape a visual loom immediately when already pursuing prey. Instead, mice of both ages (adolescent and adult) and sexes reliably froze first upon loom onset. Following this stereotyped initial freeze, subsequent orienting choices were highly variable: whether mice only froze, escaped to shelter, or resumed pursuing the sweeping stimulus (other) depended on sex, age, and the age at which they had gained rewarding prey-hunting experience. Notably, every group tested showed evasive choices or dynamics that were uniquely shaped by rewarding hunting experience at specific ages. However, observed changes in orienting overall could be classified as either down-regulating or up-regulating shelter return (escape) behavior. This is an important replication and extension of prior studies showing visually-evoked sequencing of freeze and escape behavior (Shang et al., 2018), sex-dependent evasive action choice (Shang et al., 2015) and experience-dependent flexibility in loom-evoked escape behavior (Lenzi et al., 2022).

Overall, we find meaningful variability in evasive action choice that depends on age, sex, and the developmental timing of stimulus salience learning. This work suggests distinct, potentially conserved evolutionary payoffs for different choice types across ages and sexes. Because predation risk is highest while animals are engaged in survival-critical behaviors like foraging and socializing (Lima & Dill, 1990; Salido et al., 2023), our paradigm offers a more realistic way to study visual stimulus prioritization. It revealed a robust, consistent initial loom response across all groups—disengagement from ongoing visual processing (reduced locomotion speed and prey-localization accuracy)—followed by a more variable, delayed shift in probability of returning to shelter. These findings identify previously unrecognized factors shaping visual orienting decisions within a single species. Crucially, this was discovered within a mammalian species where we are armed with knowledge of which with brain regions and circuit connections may be most likely to be altered to induce the observed changes, changes downstream of the superior colliculus for example (Shang et al., 2015 and Li et al., 2023). Thus, this work has strong implications for enhancing our mechanistic understanding of failures in stimulus prioritization in natural environments, predicting prey choice and predator success in the wild, and identifying developmental mechanisms that may be evolutionarily targeted to stabilize behavioral outcomes and minimize predation risk across species.

## Results

### Evasive action choice and dynamics in a visually-driven approach-avoidance conflict assay are sex and age dependent in mice

We developed a visual loom presentation protocol for mice that is triggered by their approach to visual objects of varying appetitive salience (**Fig. 1A and Supplemental Fig. 1**). We first habituated naive mice of both sexes and two age groups to our previously developed behavior arena (Procacci et al., 2020) with an added shelter. We then presented sweeping visual motion stimuli from a computer screen near the bottom of the arena. Three speeds of sweeping motion stimuli as in Proccacci et al., 2020 were presented to the mice for 5 minutes each. Mice and visual stimuli were tracked with Deeplabcut Live (Nath et al. 2019; Kane et al. 2020) and measures of mouse speed, range from target, and bearing of the mouse relative to the sweeping stimuli (stimulus angle) were generated in real-time. When the conditions were met that the mouse was within 5 cm of the sweeping visual stimulus, averaging a locomotor speed of at least 15cm/sec and facing the target within 90 degrees of head-cantered azimuth (**Fig. 1A**), a highly-salient visual loom stimulus, consisting of a train of three consecutive looms, was triggered above the mouse’s head. The loom stimulus was a rapidly expanding dark contrast disk presented three times in succession with an interstimulus interval of 0.1 seconds lasting overall 1.8 seconds total. The disk expanded from 0 to 20 degrees in 500 milliseconds similar to prior studies of visual loom only responses in mice which instead were triggered based on a mouse’s entry into the middle of an arena (Shang et al., 2018). Under these conditions, we compared basic features of locomotion and exploratory behavior (**Supplemental Fig. 2**) as well as stimulus locked responses (**Fig. 1 & Fig. 2**). Some mice failed to leave the shelter for prolonged periods and approach the sweeping stimulus even after 15 minutes and were thus eliminated from competing stimulus presentation analysis. However, there were no differences in the percentage of such mice that depended on age or sex (**Supplemental Fig. 1A** ethograms), nor were there significant differences in mean “time spent in shelter” between naïve experimental groups (**Supplemental Fig. 2B**). For each mouse, after habituation to the arena, sweeping stimuli were presented and most subjects in all groups of mice approached the sweeping stimulus speeds that typically evokes the most innate approaches in adult and adolescent mice of both sexes, 2 and 15cm/sec (Proccaci et al. 2020; and Gonzalez-Olvera et al. 2025).

**Figure 1:**
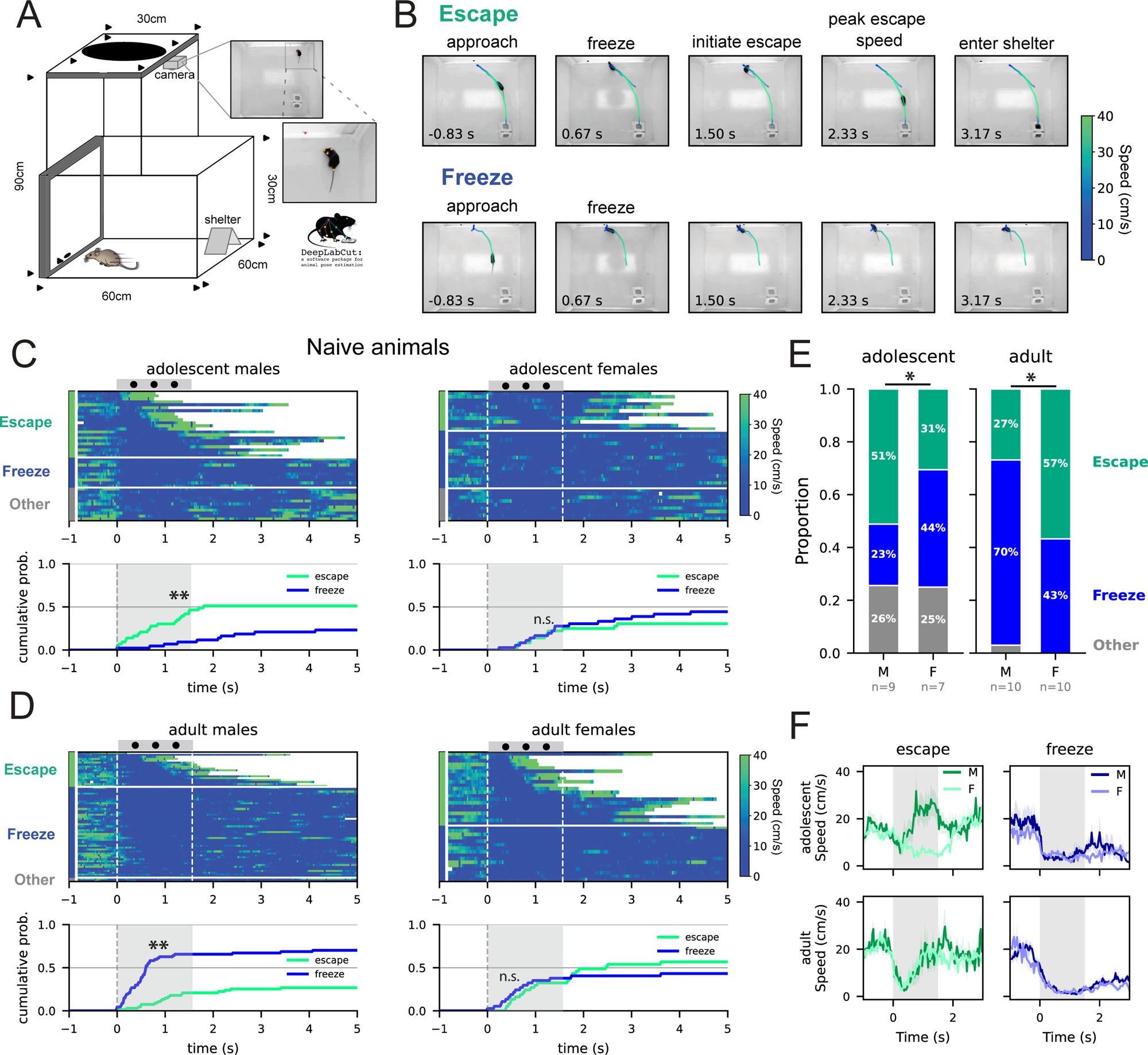
A competing valence visual stimulus paradigm that mimics prey foraging - predator avoidance conflict reveals sexually dimorphic differences in trade-off strategies in mice. (A) Closed-loop paradigm used in our experiment, as the mouse chases the visual target, an overhead loom is triggered. Mouse position is tracked in real time using DeepLabCut live. (B) The two most frequent orienting responses to the overhead loom conflicting with lower sweeping stimulus: freeze + escape defined as ‘escape’ and prolonged freezing without escape defined as ‘freeze’. (C, D) Ethograms representing mouse locomotion speed aligned to first loom onset for mice across different age and sex. Each row represents the locomotion speed over time in a single trial. The left most column of the ethogram represents if a trial was classified as escape, freeze or other. Dashed dotted lines represent looming stimulus onset and offset. Plots below each ethogram represent the cumulative probability of each group escaping or freezing only in response to the looming stimulus. (E) Proportion of trials where mice for each age and sex escape, freeze or display other behavior related to continued pursuit of sweeping stimulus. (F) Loom onset aligned average locomotor speed across trials for mice from each age and sex. Darker lines represent average across male animals and lighter lines across females. Shaded bands around the mean line represent the SEM. Cumulative probability data analysed by Kolmogorov-Smirnov test, *= p < 0.05, **= p < 0.01, ***= p < 0.001, escapes: N = 22,11,18 & 21 and freezes: N = 10,16,47 & 16, adolescent males, adolescent females, adult males versus adult females, respectively. Proportion data analysed by Pearson’s Chi-square test with yates correction within age groups, *= p<0.05, **= p<0.01.

**Figure 2:**
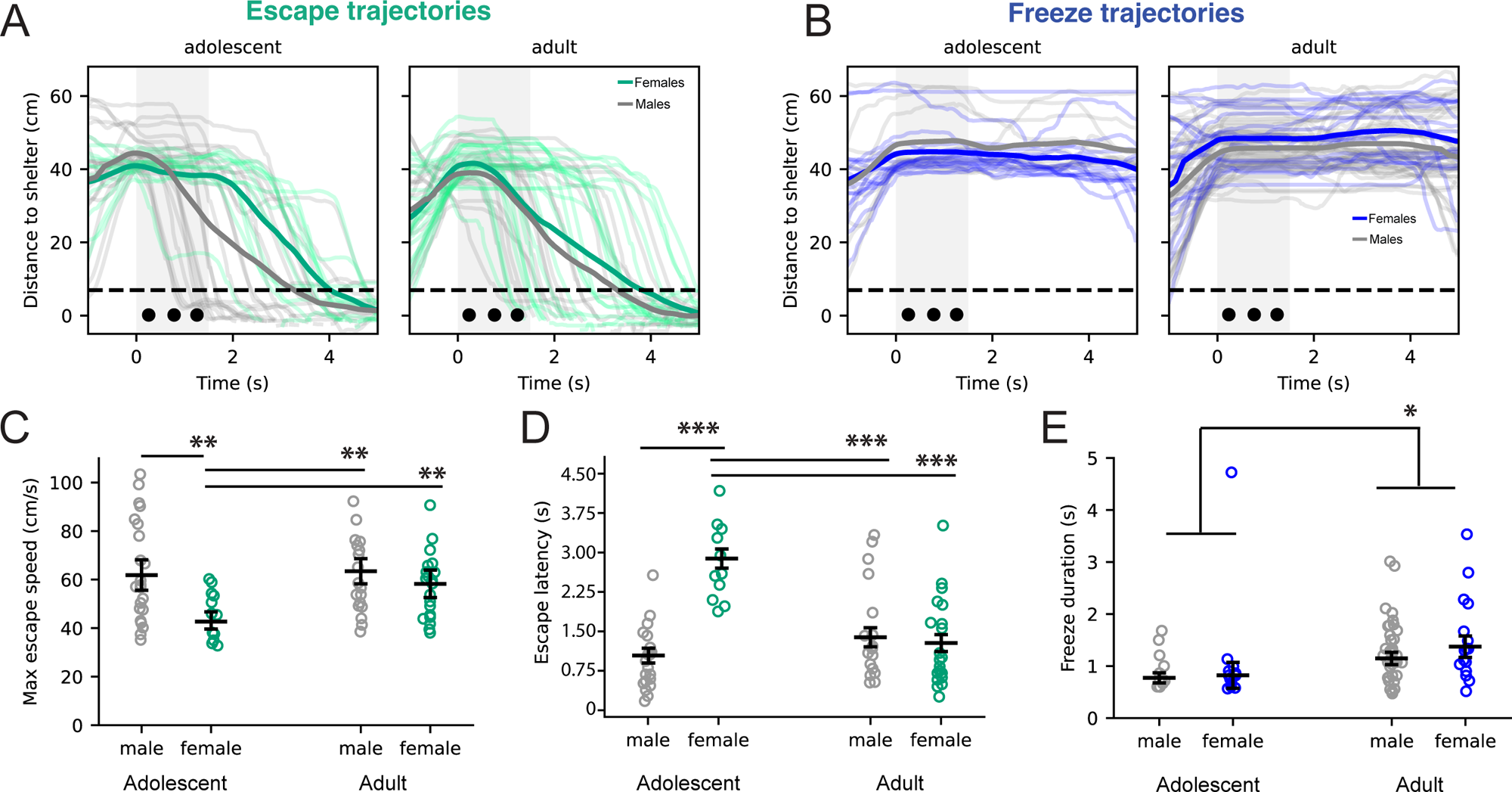
Escape dynamics are sexually dimorphic specifically during adolescence, (A) escape trajectories and (B) freeze trajectories relative to distance from the provided shelter. The dark lines represent mean across aggregated trials for the specified group. (C) Means for maximum escape speed, (D) escape latency and (E) freeze duration across age and sex for naive mice. Error bars represent standard error of mean (SEM). Each trace and data point represents a single trial. No main effect of subject was found. Data analysed by mixed effects, unbalanced, Two-way ANOVA, followed by Tukey’s HSD posthoc testing, sex and age as independent factors, *= p < 0.05, **= p < 0.01, escapes: N = 22,11,18 & 21 and freezes: N =10,16,47 & 16, adolescent males, adolescent females, adult males versus adult females, respectively.

Mice across all groups exhibited three mutually exclusive categories of measurable responses to loom in this context: 1) period of immobility only, lasting 500ms or longer, labeled as “freeze”, 2) a freeze (> 500ms) or arrest (immobility between 100 and 500 ms) followed by a run to shelter labeled as, “escape” or, 3) “other” which contains variable responses related to continued sweeping stimulus engagement or remaining near the sweeping stimulus (**Fig.1B and supplemental video 1**). Freezes were formally classified as periods of immobility of greater than 500ms as in prior studies (Yilmaz et al., 2013; Shang et al., 2015; Shang et al., 2018; Evans et al. 2018; Lenzi et al. 2022). Escapes in the context of our approach-avoidance conflict visual stimulus paradigm were notably characterised by significant reductions in locomotor speeds prior to escape in nearly all subject types (**Fig. 1B & F**), but varied between 0.1 and 1.5 seconds long. In prior research, pauses in locomotion under 500ms have been varyingly referred to as arrests or freezes and often referred to as freezes when produced in a context that was inferred to be aversive. Here, we operationally define pauses longer than 500ms in response to loom as freezes as these could be also observed in the absence of escapes (run to shelter, Shang et al., 2015 and Shang et., 2018). In addition, we also directly show the variability in modulation of locomotion behavior over time relative to stimulus onset to better facilitate objective cross study comparisons based on this as a continuous measure. Regardless, the categorical classification of pauses in movement for greater than 500 ms in the context of an overhead loom stimulus as freezes in this study will allow more direct comparisons of past work in this area.

Adolescent males were significantly more likely to escape to shelter following the loom stimulus rather than freeze relative to adolescent females (**Fig. 1C**, p = 0.00124, KS-test, and **Fig. 1E**, left, p = 0.0495, Chi-square test, N=9 v. 7, male v. female, respectively). In adult mice, the reverse trend for sex was observed, with adult females ultimately escaping more often than freezing only relative to adult males (**Fig 1D**, p = 0.00003289, KS test and **Fig. 1E**, right, p = 0.0433, Chi-square test, N= 10 v. 10, male v. female, respectively). This replicates early findings for sexually-dimorphic responses in adult mice in response to loom (Shang et al., 2015). Thus, following an initial dip in locomotor speed with the first appearance of an overhead loom, 51% of adolescent males and 57% of adult females, versus 31% of adolescent females and only 27% of adult males, choose to escape to a provided shelter after overhead loom when already fixated on a sweeping target (**Fig. 1E**).

Related, adolescent males were the least likely to freeze for durations greater than 500ms (23% vs. 44%, 70% and 43% for adolescent males, adolescent females, adult males and adult females, respectively). Altogether, attending and pursuing a visual object of interest leads to highly stereotyped freezing responses to visual loom presented in the peripheral, overhead visual field in mice regardless of sex and age. However, the subsequent choice to escape to a shelter following the initial stimulus-induced freezes are variable and show a robust sex by age interaction. The response of adult mice in this context is similar and consistent with studies stimulating loom-responsive circuits in the superficial and medial superior colliculus (sSC) directly (Shang et al., 2015). This region of the SC encodes binocular input.

In addition to scoring overt changes in orienting based on loom, we further quantified the dynamics of escape and freeze behaviors as distinct neural circuits downstream of the SC are known to regulate these aspects of the behavior versus the choice of whether or not to freeze or escape itself. We quantified and found significant differences in escape latency, max escape speed and freeze duration that were dependent on an age by sex interaction (**Fig. 2**). Female adolescent mice were significantly slower during an escape (**Fig. 2A & C**, p = 0.03121, main effect of sex by age interaction, unbalanced, Two-way ANOVA, and p = 0.0058392, adolescent females versus adolescent males, Tukey’s posthoc correction, N = 22, 11, 18 & 21, adolescent males, adolescent females, adult males and adult females, respectively). Adolescent females also took significantly longer to initiate an escape relative to all other groups which were not significantly different from each other (**Fig. 2A & D**, p= 1.387^^10-6^, main effect of sex by age interaction, unbalanced, Two-way ANOVA, and p= 5.067^^10-9^, adolescent females versus adolescent males, Tukey’s posthoc correction, N = 22, 11, 18 & 21, adolescent males, adolescent females, adult males and adult females, respectively). We found no significant differences in freeze durations by sex at each age, however, there was a main effect of age only with adults freezing significantly longer (**Fig. 2B & E**, p = 0.0494, main effect of age, unbalanced, Two-way ANOVA, p =0.0457, adolescent mice versus adult mice, Tukey’s posthoc correction, N = 26 and 62, adolescent freezes versus adult freezes, respectively). These results taken together demonstrates that in addition to sex and age specific difference in evasive action choice, the dynamics of those choices are also significantly different and depend on age by sex interactions (escape dynamics) or by age alone (freeze durations). In particular, adolescent female mice are least prone to escape and return to a given shelter most slowly which combined could indicate reduced aversive weighting of loom, or enhanced appetitiveness of the sweeping stimulus, or lack of ability to rapidly switch states. Any of these interesting possible explanations are not mutually exclusive. The presence of a distinct sexual dimorphism in adults is also intriguing and supports the idea that mice require flexible weighting of visual threat responses that are related to their distinct ecological needs across life.

### Prey hunting experience differentially alters evasive action choice and dynamics

Our experiments complement and expand our understanding of existing observations of the maturation of escape responses evoked by visual loom stimuli in mice. We validated predictions garnered from other species such as flies, fish and frogs, that mice will innately prioritize appropriate responses to overhead loom threat even when engaged with their own visual pursuit behaviors. However, in nature, there may be conditions where this prioritization shifts naturally such as during adolescence when mice are known to de-weight threats and engage in riskier behaviors (Gerhard et al., 2021). Alternatively, animals may bias competition of visual resources towards appetitive stimuli if they are sufficiently rewarding. Moreover, both prey pursuit and the reliable processing of overhead threats in mice have been shown to rely on binocular processing in overlapping regions of the upper binocular visual field (Johnson et al., 2021; Broersen et al., 2025; Wallace et al., 2013.) This suggests that mice could be more vulnerable to overhead threat if engaged in pursuit of visual stimuli that they learned represented a high level of reward such as insect prey.

To test how rewarding experience with target stimulus information impacts avoidance behavior, we re-measured approach-triggered loom responses in adult mice that gained successful insect hunting experience at different developmental stages, adolescence (early), P35-P45, versus 2.5 months old (late) (**Fig. 3A and Supplemental Figures 1C & D**). Insect hunting experience significantly enhances sweeping motion approaches and is a rapid and ethological way to enhance visual motion stimulus salience and value in mice allowing developmental study of rapidly induced plasticity in visual orienting (**Supplemental Figure 1C & D**, **Supplemental Figure 2A,** Allen et al., 2022; Johnson et al., 2021). Consistent with previous studies we found that independent of age of hunting experience, all experienced mice had significantly more approaches to the visual target that triggered the loom than naive mice (p < 0.01, main effect of experience, balanced, Two-way ANOVA, N = 8 subjects in each group, **Supplemental Figure 1C & Supplemental Figure 2A,** Procacci et al., 2020). We also measured approach behavior in a separate cohort of mice with hunting experience at either adolescence or adulthood where no loom were presented (**Supplemental Figure 3**). Approach number to our sweeping stimuli in the absence of any aversive cues were particularly increased in females that had hunting experience as adolescents (**Supplemental Figure 3A & B**). Additionally, all experienced mice tested in the visual conflict assay approached the visual target more accurately than mice without hunting experience regardless of age or sex (**Supplemental Figure 4,** mean = 18.12 degrees, SEM = 3.89 degrees versus mean = 38.20 degrees, SEM = 4.77 degrees, p < 0.01, Watson’s U). Thus, hunting experience generally enhanced behaviors that indicated an increased weighting of positive valence towards the sweeping stimulus (**Supplemental figures 1-3**).

**Figure 3:**
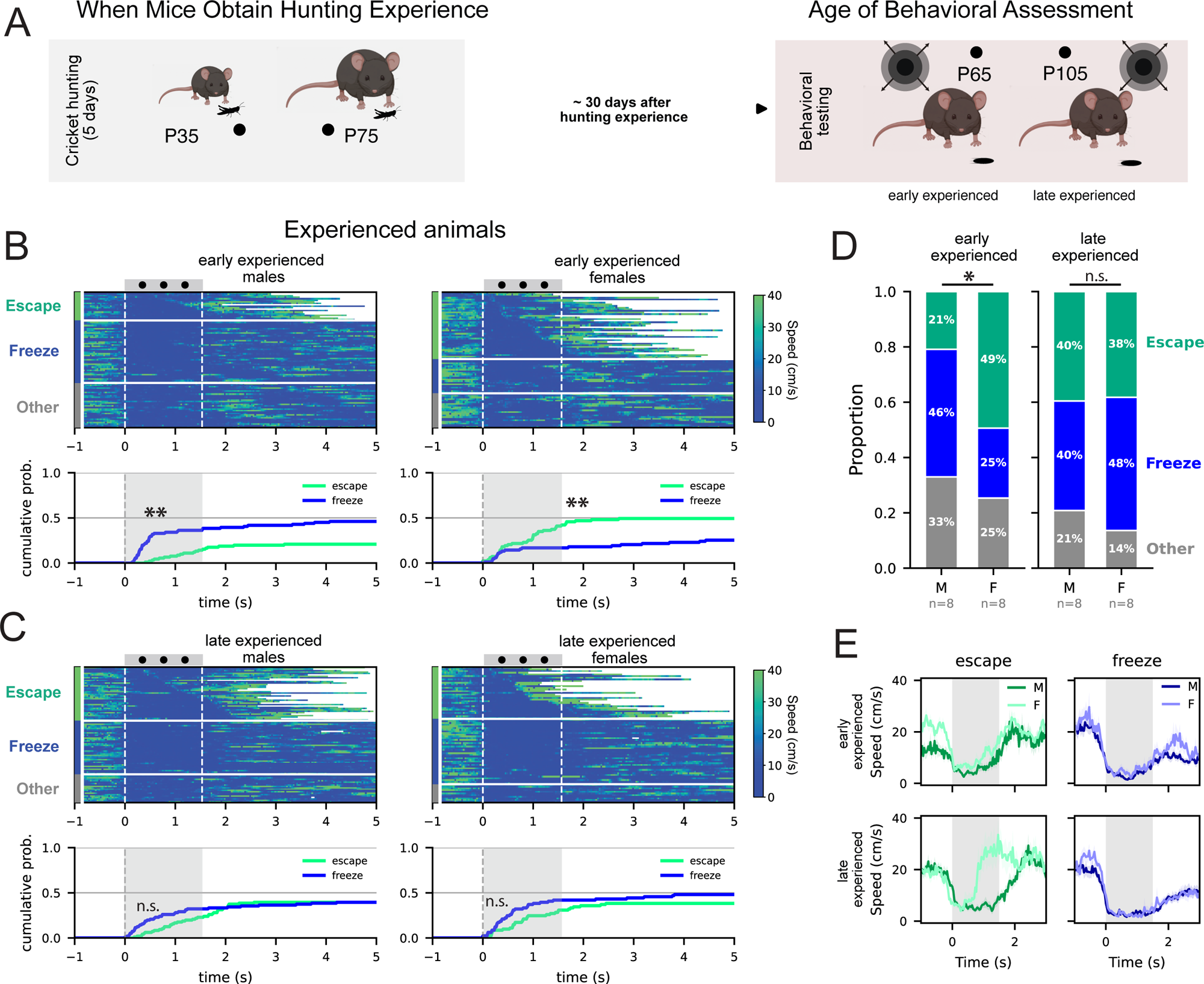
Hunting experience specifically in adolescence leads to enhanced sexual dimorphism in avoidance behavior choice. (A) Mice were provided with cricket capture experience as adolescents or adults before being tested on our competing stimuli paradigm when both groups matured into adults (B) Ethograms representing the speeds of mice with prey hunting experience as adolescents (early) relative to the triggering of overhead loom from approaching sweeping stimuli (plotted above the ethograms). The y-axis represents the individual trials, cumulative distributions of measured loom-evoked responses shown below, escapes in green, prolonged freezes only without escape in blue. Asterisks indicated significant differences, **= p < 0.01, KS test, N = 19 & 41 escapes for early experienced males and females, respectively and N = 42 & 21 for early experienced males and females, respectively. (C) Ethograms obtained from mice with hunting experience as adult (late experienced), presented similarly to data in B. n.s.= not significant, KS-test, N = 38 & 31 escapes for late experienced males and females, N = 38 & 39 freeze only for late experienced males and females, respectively. (D) Proportion of mice from both experience groups (hunting early versus hunting late), that exhibit escape (green), freeze only (blue) or continue to pursue sweeping motion (gray). *= p < 0.05, proportion data analyzed by Pearson’s Chi-square test with Yates correction, N= 8, 8, 8 & 8, early experience males, females and late experienced males, females, respectively. (E) Mean locomotion speed for each group over loom stimulus period (gray shading overlay). SEM represented by thickness of line.

The choice to escape from loom to a shelter were differentially impacted by early experience between males and females. Hunting experience at adolescence uniquely decreased escape responses in males and increased in females (**Fig. 3B & D**, cumulative distributions, **= p < 0.01, KS-test, & **Fig. 3D**, left, *= p < 0.05, Chi-square test, N = 8 in each group). In contrast, there were no significant differences in the proportion of freeze only versus escape decisions between adult male and female mice receiving prey capture experience when they were already mature (**Fig. 3C & D**). Although, experience did increase the number of approaches made by both males and females towards sweeping stimuli (**Supplemental Fig. 1 and Supplemental Figure 2A**). As a consequence, this generated an overall increase in the quantity of measured avoidance behaviors to the co-occurring loom presentations.

Interestingly, avoidance responses did not habituate fully during the 5-minute presentation period (**Supplemental Figure 1D**). Instead, we observed more escapes to concurring loom stimuli for all experienced groups except the males who received hunting experience specifically as adolescents (**Fig. 2A** vs. **4A** and **Fig. 5A** vs. **5B**, interaction between sex and age of experience, p < 0.05, balanced, Two-way ANOVA, N = 8 mice in each group). To visualize more directly how hunting experience altered the relative balancing between approach and avoidance, as both changed with appetitive hunting experience, we created a ratio between mean number of approaches to mean number of escapes per group (pie chart insets in **Fig. 5A & 5B**). The comparison of approach to escape ratio for each group demonstrated that the relative number of target approach to escapes is initially sexually dimorphic in adolescents with males skewed towards escape and females towards approach (**Fig. 5A**, inset, p < 0.01, Fischer’s exact test, N = 9 and 7, adolescent male and female, respectively). This trend is reversed in stimulus naïve adult mice, where adult male skew towards approach and females skew towards escape (**Fig. 5A**, inset, p < 0.05, Fischer’s exact test, N = 10 and 10, adult male and female, respectively). Intriguingly, hunting experience specifically during adolescence leads males to skew more heavily towards approach, with a significant increase in approach and a significant decrease in escape (**Fig. 5B**, inset **and Supplemental Fig. 1C,** ethograms). This led to a significant difference between the approach to escape ratio between early experienced male and females (**Fig. 5B**, inset, p < 0.01, Fischer’s exact test, N = 8 and 8, early experienced males and early experienced females, respectively. Instead, hunting experience generated similar relative ratios in male and females when experienced late, which indicated that the increase in approaching a moving target was offset by an increase in escape triggered by loom.

Overall, males with hunting experience displayed a unique imbalance between approach and avoidance behavior in this context. This indicates a sensitive period for adolescent males whereby naturally rewarding pursuit experiences reduce and alter threat responses when they are already engaged with rewarding stimuli. This would render this population of individuals more susceptible to predation as adults if the predator had an advantage with slower moving or freezing prey. On the other hand, the late experienced females escaped with faster escape speeds relative to all other groups (**Fig. 4C**, main effect of sex by age of experience interaction, p < 0.001, Two-way ANOVA, Tukey’s posthoc testing, **Fig. 5B**, yellow colored/high speed, escape more abundant, **Supplemental Video 2**). Females gaining hunting experience at any age also responded by more quickly mounting escape at loom onset (**Fig. 4D**). This demonstrates that unlike adolescent males, females increase the efficacy of threat-induced escape behavior to offset their increased interest in rewarding visual stimuli (**Fig. 3E & 4A, C & D**). Thus, age at which prey hunting experience occurs, and enhances the salience of the sweeping stimulus, differentially impacts evasive action choice and escape dynamics in mice in a sex-dependent manner. Our results support that increases in the perceived appetitiveness of the sweeping stimulus alters approach-avoidance conflict behaviors. Males display less aversion of the co-occurring loom stimulus by downregulating escape related behaviors. This is especially strong for males that gain positive experience with the rewarding stimuli as adolescents. Females overall modulate escape probability and dynamics to better facilitate switching between approach and avoidance orienting.

**Figure 4:**
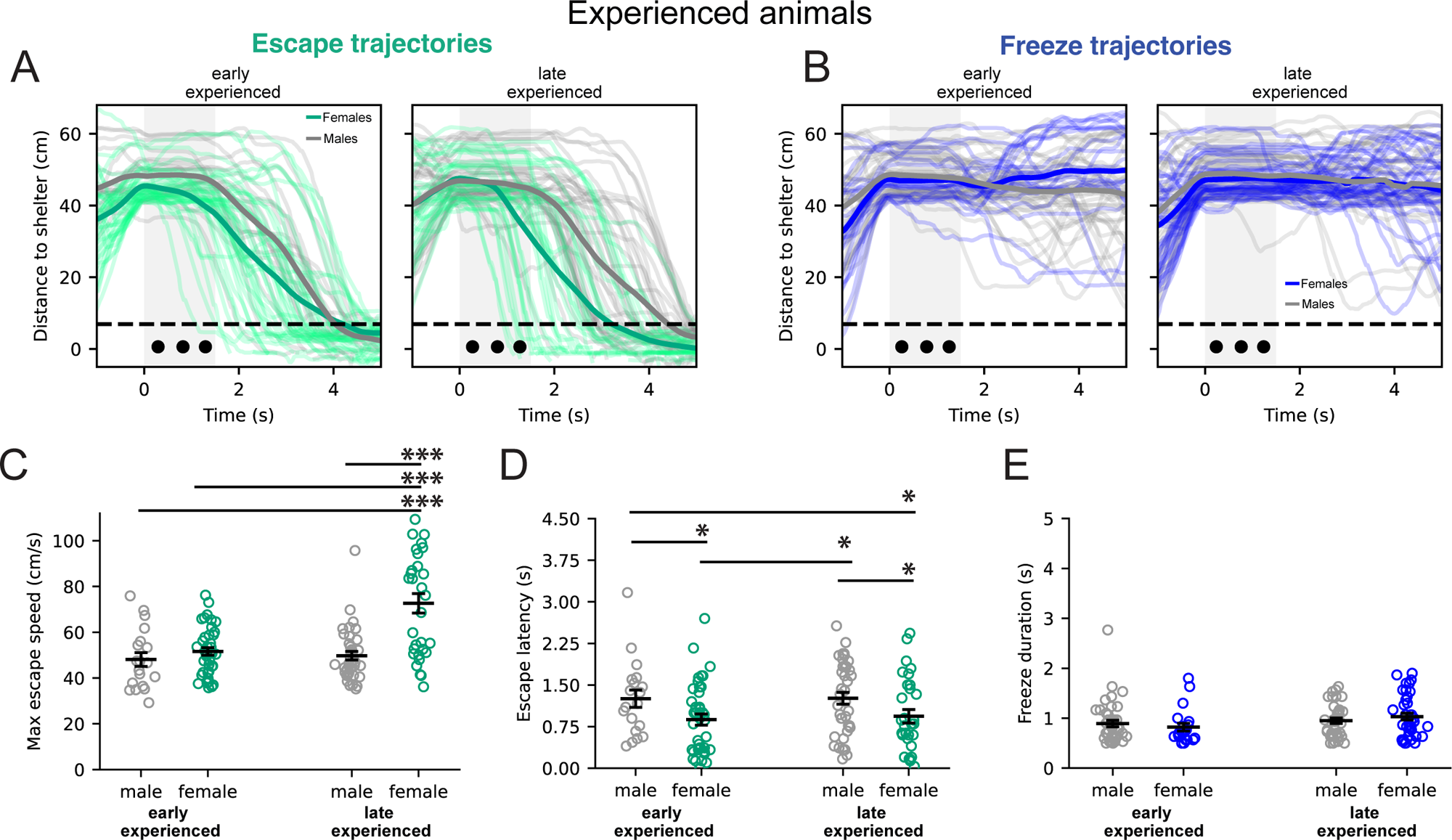
Escape dynamics can be enhanced by hunting experience in females regardless of age of experience. (A) escape trajectories and (B) freeze trajectories relative to the distance from shelter. The dark lines represent Mean across trials for the specified groups. (C) Mean maximum escape speed shows enhanced escape speed uniquely for females who hunted as adults. ***= p < 0.001, interaction between sex and experience, Two-way, unbalanced ANOVA, Tukey’s posthoc correction, N= 19, 41, 38 & 31 escapes for early experienced males, females and late experienced males, females, respectively. Error bars SEM. (D) Mean escape latency is reduced for females regardless of when they experience hunting. *= p < 0.05, main effect of sex, Two-way, unbalanced ANOVA, Tukey’s posthoc correction, N= 19, 41, 38 & 31 escapes for early experienced males, females and late experienced males, females, respectively. Error bars SEM. (E) freeze duration by age of experience and sex. No significant effect of sex, nor age of experience, nor interaction, Two-way, unbalanced ANOVA, Tukey’s posthoc correction, N= 42, 21, 38 & 39 freezes for early experienced males, females and late experienced males, females, respectively. Error bars SEM. Each data point represents a single trial.

To better understand whether hunting experience altered the perceived aversiveness of the loom itself, we examined the efficiency of loom-evoked escape trajectories as a function of experience in mice (**Fig. 5C**). We found that males who hunted as adolescents uniquely took more circuitous routes (less efficient paths) back to the shelter (**Fig. 5C**). This observation in conjunction with an overall reduced number of total escapes for this group (**Fig. 5A & B**), enhanced escape habituation (**Supplemental Figure 1D**), and relatively slower escapes speeds (**Fig. 4C**), suggest that male mice reduced threat responsiveness more generally as a function prey capture experience in adolescence. By comparing trajectories of mice running to shelter outside of the loom presentation, we found that the early-experienced males also moved significantly slower during spontaneous runs to shelter (**Supplemental Figure 5A-C**) and were uniquely unaffected by hunting experience in their overall visits to the shelter (**Supplemental Figure 5D**) whereas all other groups visited their shelters more often outside of immediate exposure to the loom. Taken together, adult males that hunt as adolescents are less likely to seek shelter generally, take more circuitous routes to shelter faced with a loom stimulus relative to the other groups and generally display less active avoidance behavior. Thus, this group of mice appears to have overall reduced avoidance behavior as a result of hunting during adolescence. This supports the idea that hunting experience for mice, depending on age and sex, can affect two different strategies to deal with approach-avoidance conflict. One, we observe a general effect on reducing visual loom stimulus aversion, adolescent-experienced males. Two, we see that stimulus-specific increases in appetitive approach can be balanced by generating more efficient switching to active escape strategies (females and older males).

**Figure 5:**
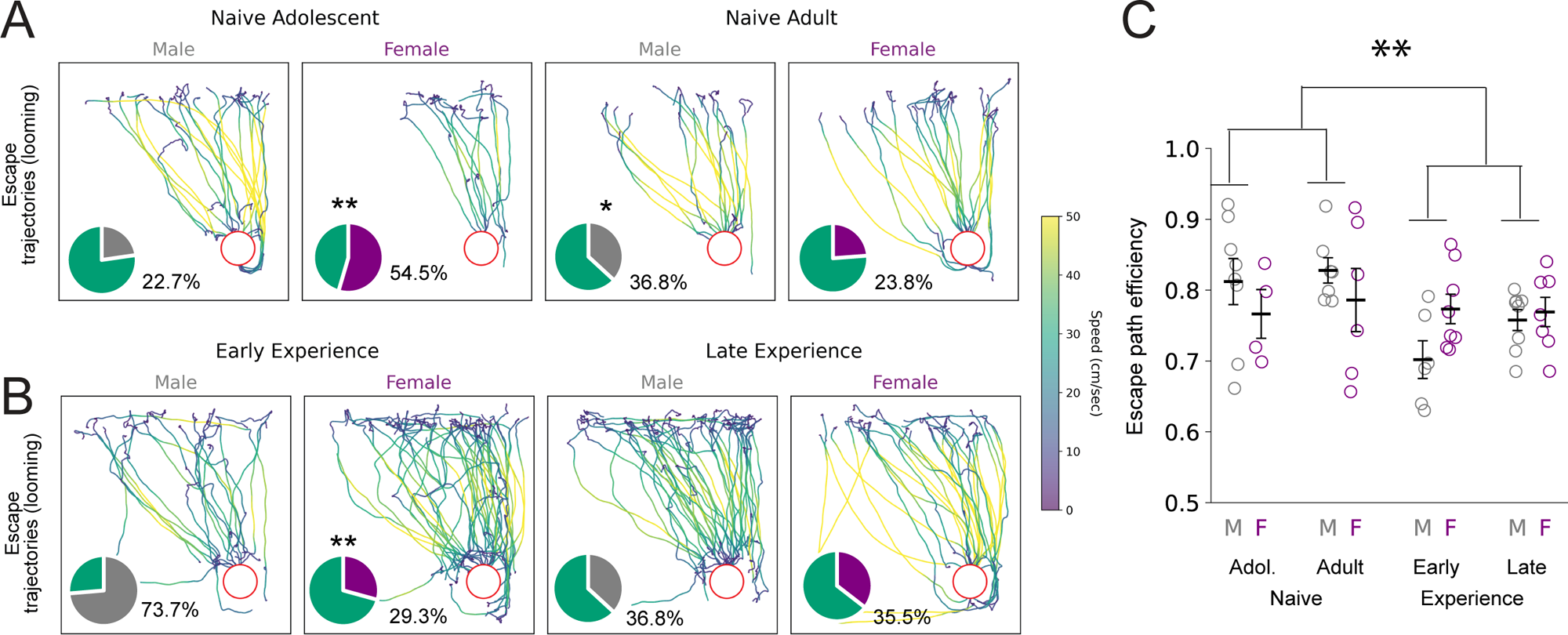
Adolescent Hunting experience uniquely shifts approach-avoidance strategy in males away from escape. (A) Mouse trajectories of naïve mice during escape trials color coded by mouse speed. Speed key, to the right, warmer colors represent faster speeds. Shelter location aligned and shown as red open circle. Pie chart insets represent the ratio of approaches to escapes for each experimental group. The ratios were calculated from the group means for approach number and escape number for each group. The gray and purple color section indicates the approach/escape ratio represented as parts of a whole. Green represents that the remaining whole is the relative escape tendency. (B) Same representations as in A except shown for developmentally timed hunting experienced mice (early versus late experience, tested as adults). (C) Mean escape path efficiency which is calculated as a ratio between the actual path length and the length of the straightest possible path between the animal’s location at beginning of loom presentation and shelter position. ** = p < 0.01, unbalanced, Three-way ANOVA, main interaction between sex and age of experience, N = 8, 4, 7,6, 6, 8, 8 & 7, naïve adolescent males, naïve adolescent females, naïve adult males, naïve adult females, early experience male, early experience female, late experience male, late experience female respectively. Purple open circles are the mean escape path efficiency measures per female producing escapes, gray open circles are the mean escape path efficiency scores per male producing escapes.

### Prey hunting experience refines the correlation between stimulus visual field location and loom-evoked orienting decisions

Finally, presenting the visual stimuli simultaneously provided the opportunity to determine whether there were finer scale correlations in evasive action choice relative to the position of the target. All loom were triggered while mice were within a certain distance from a sweeping target and pursuing that target which lead to the target appearing near centered and the loom above head. However, it is possible that the remaining variability in stimulus targeting accuracy that we saw between naïve and hunting experience mice, further explained subsequent evasive action choices. In particular, given the recent studies showing that escape and approach behaviors depend on maximizing binocular visual field overlap, we predicated the most conflict between generating immediate escape when mice were most accurately approach sweeping targets. We measured escape behavior as a function of the egocentric angle of the approached visual target. The egocentric angle was defined as the relative angle between the center of the mouse’s head position and the sweeping motion stimulus in the overhead camera (**Fig. 6A**). We compared the means of the egocentric target position at the onset of loom for trials where escape is evoked versus those with freeze only. There were no significant effects of sex on this dataset so sexes were combined. Mice who hunted during adolescence uniquely showed a correlation between the visual angle of the target stimulus and whether they escaped to the shelter (**Fig. 6B**). In this case, mice with early hunting experience only escaped if the pursued target was more peripheral at the onset of the loom. Conversely, hunting experience at any age correlated with freezes occurring when sweeping stimuli were pursued more centrally. Taken together, we demonstrate that prey hunting experience at adolescence most significantly refines the relationship between pursued visual target location and evasive action choice, likely by ensuring that animals are more engaged with the sweeping stimulus. This timing and kind of experience may therefore be a critical natural way that mice and other mammals optimize their responses in the face of competing valence visual stimuli. It will now be interesting to determine in this context whether experience alters active visual sampling of the environment in an age dependent way and/or the weighting of stimulus valence to directly control orienting in mice. Explicit studies of a predator’s success with mice performing these behaviors in the wild may further establish the ecological implication and relevance of our findings.

**Figure 6:**
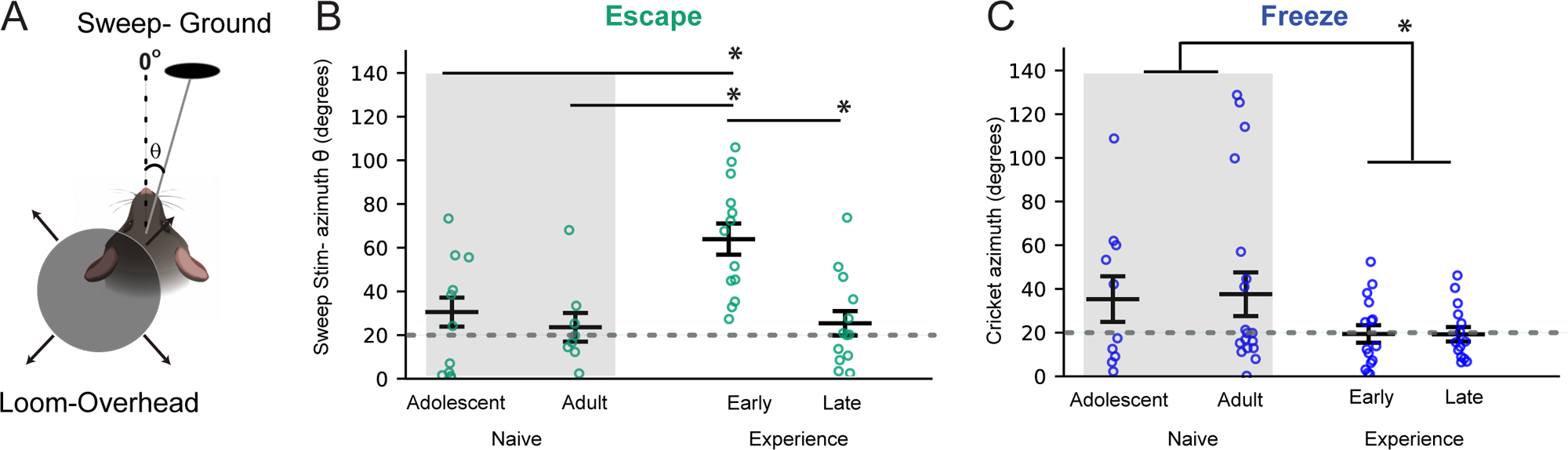
Pursued visual target location predicts escape response in mice that gained hunting experience as adolescents (early). Mean sweeping motion target angle in degrees along the azimuth at the onset of looming stimulus for different escape behavior. (A) graphic defining visual angle measurement taken at the onset of the loom start. (B) Mean sweeping stimulus angle in degrees at start of loom when mice chose to escape (green, open circles). Gray shading indicates measures from the same prey capture-naive mice shown in figures 1&2. * = p < 0.05, main effect of age X experience, N-way ANOVA, Tukey’s HSD posthoc testing, N= 11, 8, 13 & 13, naive adolescents, naive adults, early experienced versus late experienced mice, respectively. (C) Mean sweeping stimulus angle in degrees at start of loom when mice chose to only freeze (blue). Left most groups covered by gray shading represent prey hunting naïve mice shown in figures 1&2. *= p < 0.05, main effect of experience, N-way ANOVA, Tukey’s HSD posthoc testing, N= 10, 18, 15 & 14, naive adolescents, naive adults, early experienced versus late experienced mice, respectively

## Discussion

### A novel paradigm to study mechanisms of visually-guided approach-avoidance trade-offs and decision making

Our study sets up the ability to study how mice balance escape decisions elicited by overhead looming stimuli and approach towards a desired visual target. The paradigm enables precise quantification of visually guided decision-making in a natural context that encourages mice to engage in the active visual sampling and locomotion based-strategies critical to shaping the connectivity of their brains. Park et al. (2024) used a related approach in which food-deprived mice, presented with food pellets at the arena center, faced an overhead loom during feeding—pitting feeding against escape. Our paradigm instead isolates visual cue conflict itself: both target and loom have distinct spatiotemporal characteristics that elicit opposing responses (Procacci et al., 2020; Yilmaz & Meister, 2013). By timing the loom to the animal’s approach phase and distance from the target, we can also control where in the visual field the loom appears—enabling study of visual competition in a freely moving animal, previously feasible only in head-fixed or fixating preparations (Desimone & Duncan, 1995b; Reynolds et al., 1999). By centering the work on visual conditions, we will be allowed to more directly trace the timing of neural activity through the relevant brain network during freely moving and dynamic behavior. In turn, we can identify key phases and sequencing of behavior that allows us to isolate visual stimulus encoding from decision making variables and motor outputs under more natural conditions. Critically, without the need for extensive training or conditioning of mice, we can study more rapid learning processes across a greater breadth of development.

### Trade-off Decisions Depend on Retinotopic, Head-Centered Location of Competing Stimuli

Across species, the visual system is retinotopically organized and not all cell types are equally distributed across the surface. Instead, specific cell types with ecologically relevant feature extraction are localized to specific regions of the visual field (Baden et al., 2020; Sedigh-Sarvestani & Fitzpatrick, 2022). This asymmetric localization is shaped by both natural scene statistics (Simoncelli & Olshausen, 2001) and species-specific ecological needs (Qiu et al., 2021). In mice, responses to both looming and prey-like stimuli are known to be driven by binocular visual field overlap (Johnson et al., 2021; Wallace et al., 2013).

Mice hunting crickets keep prey within the region of greatest binocular field overlap, a region dense in sustained ON alpha retinal ganglion cells that may be specialized to detect prey-like or far-away sweeping objects (Holmgren et al., 2021; Oesterle et al., 2025). Because mice keep pursued targets within a specific region of the visual field, presenting the loom at a fixed position relative to the pursued target ensures that loom falls within a defined visual field region. This in turn allowed us to assess how loom and target compete for perceptual salience in driving escape decisions relative to their most probable initial retinal stimulating position. For example, only mice with adolescent hunting experience consistently escaped when the target was likely to be more in the periphery. On the other hand, mice froze or failed to respond to the loom when the target was likely binocular or peri-binocular. These results confirm that freezes may be invoked with visual stimulation more broadly across the visual field and simultaneous to ongoing approach, while escape more directly competes with approach responses. In a freely moving context where visual targeting is not as controlled, escapes versus freezes may have been driven instead by how near the binocular visual field overlap the loom stimulus landed. This could have varied by the relative overall vigilance and active visual sampling behavior differences in mice tested under different laboratory conditions and by different groups. This is so far not known for much of the past work done measuring orienting response to loom in freely moving mice. Our findings here also indicate that lack of sampling adequate numbers of each sex may also have contributed to past discrepancies in observations of loom-evoked defensive behavior in mice.

Experience hunting live prey, sharpened visual motion targeting in all groups. Yet, only the mice with hunting experience in adolescence showed a clear refinement of the relationship between pursued target location in the azimuth and escape response evoked by overhead loom. There may be two possible explanations to account for this. First, binocular receptive field properties may shift with hunting experience only in adolescence. Bissen et al., 2026, found that temporal frequency discrimination improves in adolescent mice hunting during P28–P35. If hunting experience shifts the center-surround location of binocular receptive fields in the SC—as this improved discrimination would suggest—the relative positions of loom and target with respect to that center-surround structure would change, altering the response to the competing stimuli. This could reflect divisive normalization of target and competitor responses, or a winner-takes-all mechanism as seen in the SC for competing stimuli in head-fixed mice (Banerjee et al., 2025). This possibility assumes that the two stimulus types mice were concurrently processing overlapped spatially in where they stimulated the retina, e.g., near the binocular visual field. A second possibility is that eye-head coordination itself changes substantially with adolescent cricket-capture experience only. Such differences could shift where the loom stimulus falls in the visual field, driving the difference in escape responses. Testing this would require eye tracking coupled to performance in this assay. The overall absence of escapes when adolescent hunting-experienced mice are accurately pursuing stimuli compellingly supports the idea that engaged binocular processing of prey-like stimuli interferes with active avoidance of predation. This in turn is consistent with studies that show that mice alter head position, down or up, during either prey hunting or to escape from overhead predators to process the most salient information in the region of best binocular visual field overlap. However, freezes only will still be elicited if loom hit the upper visual field while mice are pursuing accurately targets. Indeed, we observed characteristic up-ward shifts in head position upon loom appearance that disrupted sweeping targeted pursuit (**Supplemental Video 1**). Thus, our data are consistent with studies that show that visually-guided escape choice is both dependent upon visual field location of the stimulus, near or in the binocular visual field, as well as downstream gating mechanisms that route that information to different motor outputs (Broersen et al., 2025; Shang et al., 2018; Li et al., 2023).

### Trade-off Decisions Depend on Intrinsic, Life-History-Dependent Features of Mice

Responses to this paradigm varied by age, sex, and age of gaining hunting experience, with both experienced and naive mice showing distinct patterns when facing competing appetitive and threatening stimuli. Naive adolescent males were more likely than adolescent females to escape, particularly on first exposure to the loom during pursuit—potentially reducing their vulnerability to predation at this stage. However, after this first exposure, adolescent males rapidly habituated and became less likely to escape than other groups, consistent with other work studying the ontogeny of avoidance response to loom stimuli alone (Albrecht et al., 2025). Notably, adolescent males with prey-hunting experience went on to de-weight risk and dampen escape behavior as adults—a shift that would increase predation vulnerability. We speculate that mice pursuing a highly salient target are most likely to be facing it directly and freeze at loom onset, heightening vulnerability. This is consistent with the idea that hyperfixation on reward carries a cost in the presence of predators. Females appear to counter this vulnerability differently: after prey-capture experience raises target value, they either escape more frequently or escape faster from loom onset. The two sexes thus appear to pursue distinct strategies in this approach-avoidance conflict, differently shaped by experience-driven changes in target value.

Our findings are reminiscent of early observations in mice of sex-divergent strategies in responding to threat under the fear-conditioning context (Gruene et al., 2015) and may indeed involves shared circuit underpinnings. Escape and freeze responses to overhead looms are thought to serve distinct adaptive functions against aerial predators: fleeing may favor dense foliage, while freezing may better evade detection in open habitats by disrupting motion cues. Baier et al. (2025) showed that two closely related deer mouse species diverge in exactly this way—freezing versus fleeing—reflecting adaptation to dense versus open habitats. Why sexes and ages within a single species adopt distinct strategies is less clear; one hypothesis is that dispersal, foraging, and mate-seeking differ by sex at the ages we examined. Studies of *Mus musculus* under rewilded or more naturalistic conditions may help clarify this understudied aspect of mouse ecology. Regardless, we speculate that the circuits underling this evolutionarily-shaped approach-avoidance conflict process may be particularly relevant to previously described studies of a sensitive period for optimizing approach avoidance behaviors generally in adolescent mammals (Gerhard et al., 2021).

Beyond environmental influences, sex and developmental stage shape the general weighting of reward against risk. Naive adult mice froze significantly more often than adolescents, consistent with Albrecht et al. (2025), who found that mature glutamatergic (GluA2-containing) synapses in the dorsal PAG—a region governing both approach and escape—decrease at P30 before stabilizing. This structural change may explain why escape responses peak at P30 and decline into adulthood, and a similar mechanism could underlie the age-related pattern we observe in our conflict paradigm. Thus, the circuits that transform retinotopically-mapped stimuli into downstream motor responses also contribute to differences observed here. Thus, overall, this paradigm allows for accounting for both intrinsic factors of mice such as sex, hormones or hunger state from the concrete features of stimulus feature mapping in sensory systems, e.g. retinotopic location, stimulus speed and the interaction between each.

## Supporting information

Behavior Scoring Definitions

Escape behavior after timed experience

## Acknowledgements

We would like to thank Kelsey Allen, Kierian Huang and Aja McDonogh for their inputs and feedback on the manuscript. This work was funded by RO1 EY032101 awarded to Dr. Hoy.

## Methods

### Animals

All experimental procedures were conducted in accordance with protocols approved by the University of Nevada, Reno Institutional Animal Care and Use Committee. This study used 70 C57BL/6 mice, evenly distributed between males and females. Our groups included both adolescent (P30-45) and mature adult (P60+) mice. Initially, we used 38 mice without hunting experience (9 adolescent males, 9 adolescent females, 10 adult males, 10 adult females) to quantify the behavioral responses to our paradigm. Next, to evaluate the role of hunting experience in reweighting escape decisions based on age of experience, we collected data during our paradigm from mice that had undergone hunting experience as adolescents (8 each for males and females) or as mature adults (8 each for males and females).

Animals were group-housed in same-sex cohorts of 3-5 individuals per cage in an on-campus vivarium. Except when noted, standard housing conditions included ad libitum access to water and food (Envigo Teklad diet 2919). The facility maintained a 12-hour light/dark cycle, with all behavioral testing conducted within 3 hours following the dark-to-light transition. Food restriction was implemented only when mice were provided with cricket-hunting experience (see below) and involved removing food hoppers at the onset of the dark phase. Following this, animals were tested 12-16 hours post-restriction, immediately following the subsequent dark-to-light transition. At the end of behavioral testing, mice were returned to their cages with restored ad libitum food access throughout the light phase until the next restriction period.

### Behavior

We performed behavior in a specialized, lightproof, double-walled, sound attenuating chamber to ensure controlled environmental conditions. The testing arena consisted of a square acrylic open field (60 × 60 × 30 cm) with white vinyl flooring. The arena configuration included two opposing white acrylic walls and two opposing walls formed by Hewlett Packard VH240a video monitors (60.5 cm diagonal, 1920×1080, 60 Hz, 250 cd/m²). A third monitor was placed overhead at a height of 90cm with one of its edges aligned with the lower monitor (Figure 1A). Behavior recordings were captured using an overhead-mounted Logitech HD Pro Webcam C920 at 30 frames per second.

### All raw video files can be accessed from the available repository

https://osf.io/ufjsd/overview?view_only=3dfac5325c9e43448c5e51f268059c8d

### Hunting experience

Mice were provided hunting experience as described previously (Hoy et al. 2016, Procacci et al. 2020). Briefly, following a handling and habituation period, mice were placed individually into the arena containing a live cricket for live prey capture experience.

Hunting proficiency was considered established when food-restricted mice consistently exhibited their most efficient prey capture behavior, typically capturing the cricket within 30 seconds of its introduction into the arena.

### Visual stimuli

Computer-generated visual stimuli were created using the MATLAB Psychophysics Toolbox (Brainard & Vision, 1997) and presented on an LCD monitor (60 Hz, 50 cd/m²) in a dark room. The visual target was created to simulate prey-like characteristics, similar to stimuli that our lab has previously shown are readily approached by mice (Allen et al., 2022; Gonzalez-Olvera et al., 2025; Procacci et al.,2020) The target was programmed to sweep horizontally across the monitor at a consistent height of 2.5 cm above the arena floor.

For the overhead looming stimulus we presented three successive rapidly expanding disks. The disks expanded linearly from 0 to 20 degrees visual angle in 0.5 seconds, similar visual stimuli have been used to evoke robust escape responses (Shang et al., 2018). The interstimulus interval was 0.1 seconds.

### Competing stimulus paradigm

For the main experiment, mice were placed in the arena, which contained a shelter at the opposite corner of the screen displaying the visual target (Figure 3.1A). The arena was cleaned with isopropyl alcohol and shelter was swapped before behavior from a new mouse was collected. The mice were allowed to explore the arena for 2 minutes without any visual stimulus. At the start of the experimental trial, the visual target was displayed moving out of the corner of the screen and sweeping across the length of the screen. The target was presented at three different speeds (2, 15 and 50 cm/s) for 5 minutes each, in pseudorandom order across mice and then aligned by order of increasing speed sessions for presentation purposes in all displayed ethograms. DeepLabCut Live (Kane et al., 2020) was used to estimate the mouse’s position in real time. We tracked four keypoints on the mouse. One each on the mouse nose, left and right ear and the base of the tail respectively (**Figure 1A**). We used a custom script in MATLAB and python to project the sweeping visual target onto the overhead camera reference frame. This allowed DeepLabCut to interact with the stimulus presentation and trigger the looming response if a set of pre-determined conditions were satisfied. During an active approach to the target, we triggered a loom stimulus when the following conditions were met: the mouse was less than 5 cm from the target, moving at more than 15 cm/s, and the target was within +/- 90 degrees in its field of view. Because of a lag between sensing this conditions and translating to loom presentation, there was some variability in distance and visual angle of the sweeping stimulus at the moment of true loom presentation. We thus presented visual angle measures and estimated distance from sweeping stimulus at the time of actual loom start and aligned all mouse behaviors to this real-time onset. We also negated approaches made along the stimulus presentation monitor by ensuring that the approach analyzed initially started at a distance greater than 10 cm from it.

### Quantification of escape behavior and escape dynamics

A response was counted as an escape if the mouse returned to its shelter within five seconds after loom onset, and at some point during those five seconds, it reached a speed of at least 25 cm/s while moving toward the shelter. Even spontaneous returns to the shelter not associated with loom, saw all mice moving greater than 25cm/sec. Freezing occurred when the tracked point position did not change by more than 0.5 cm for at least 15 sequential frames ∼500ms. This definition yielded the same events as those tagged when isolating epochs where mouse locomotion speeds were less than 2.5 cm/s for greater at least 500 ms. An additional criterion for labelling a response as a freeze is that an escape to the shelter did not happen within 5 seconds from loom onset. Any movement pattern that didn’t meet the criteria for either an escape or a freeze was classified as a “other” where the mouse briefly disengaged the sweeping stimulus, but not enough to constitute an escape, and was not immobile enough to be classified as a freeze. Many of these events correlated with a resumed approach towards the sweeping stimulus if it remained on the screen.

Spontaneous escapes to the shelter were defined by searching for trajectories that started and ended outside any loom window (starting one second before loom onset to five seconds after the loom). We analyzed only those spontaneous escapes to shelter where the mice travelled a distance of at least 40cm in five seconds.

For defining the escape dynamics, the escape latency was defined as the time elapsed from when the looming was presented until the mouse began its initial movement back toward shelter, with a speed greater than 10 cm/s. For both loom-associated escapes and spontaneous returns to shelter, the maximum escape speed is the fastest speed that the mouse achieved while heading back to shelter during the five-second window after the loom onset. The spatial efficiency of escape was defined as the total distance covered by the mouse during an escape starting from the loom onset to the moment the mouse reached the shelter normalized by the shortest distance between mouse position at loom onset and shelter entrance (Claudi et al., 2022).

The pursued visual target location at loom onset was calculated by estimating the angle between the line joining the centre of the mouse head and nose, and the line joining the centre of the mouse head and the visual target (**Figure 6A**, Allen et al., 2022).

### Statistical Analysis

Statistics were performed using Python, GraphPad Prism or JASP. Means and standard error of the mean are reported in all cases except where noted in the figure legends and for target azimuthal angles where we report the circular mean. We analyzed parametric data by Two-way ANOVA or Three-Way ANOVA followed by Tukey’s HSD posthoc testing.

Distribution differences were detected via Kolmogorov-Smirnov testing. For analysis to detect difference in habituation to stimuli, and/or changes across subsequent loom presentations within subject, we used a repeated measures, Two-way ANOVA. In most cases where N’s were unbalanced, we confirmed findings using a generalised linear mixed effects models (GLMM) for statistical tests with mouse ID as a random effects factor.

Probabilities were modeled with a binomial family with a logit link function while continuous data was modeled using a Gaussian family with an identity link. We compared specific conditions wheter main effects were detected using post-hoc tests with p-values corrected using Tukey’s HSD method. Test results with a p-value of < 0.05 were considered significant, but we noted where p-values were below 0.01 and 0.001, * = p < 0.05, ** = p < 0.01 and *** = p < 0.001 in the figures themselves and/or reported exact p-values in text for data not shown in figures.

## Notes

https://osf.io/ufjsd/overview?view_only=3dfac5325c9e43448c5e51f268059c8d

### Competing Interest Statement

The authors have declared no competing interest.

