## Supplemental Figures for "GO, NO-GO: A SENSITIVE PERIOD FOR DEVELOPING SEX-SPECIFIC VISUALLY-GUIDED PREY PURSUIT-PREDATOR AVOIDANCE TRADE-OFF STRATEGIES"

### SUPPLEMENTAL FIGURE 1

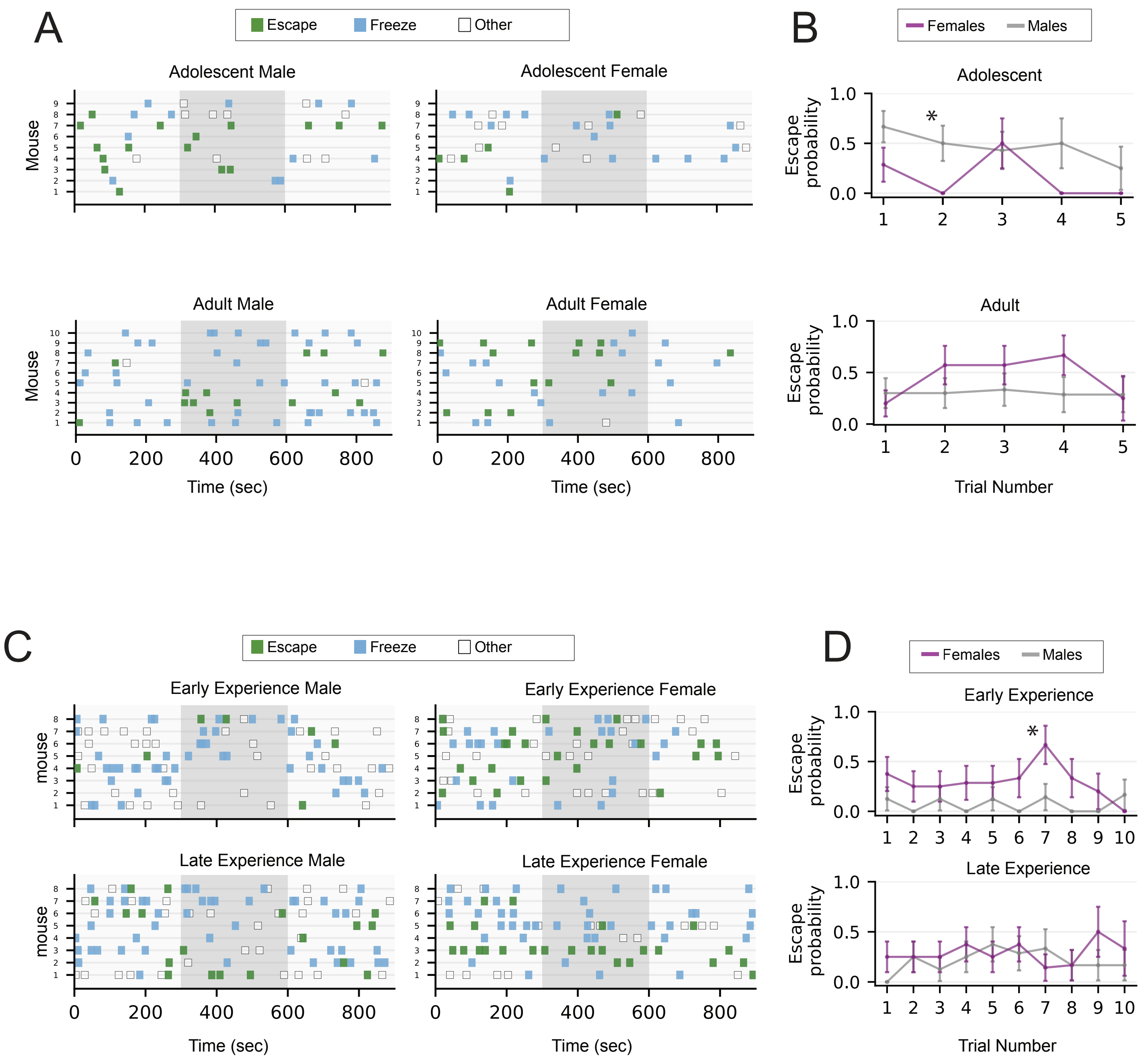

### SUPPLEMENTAL FIGURE 2

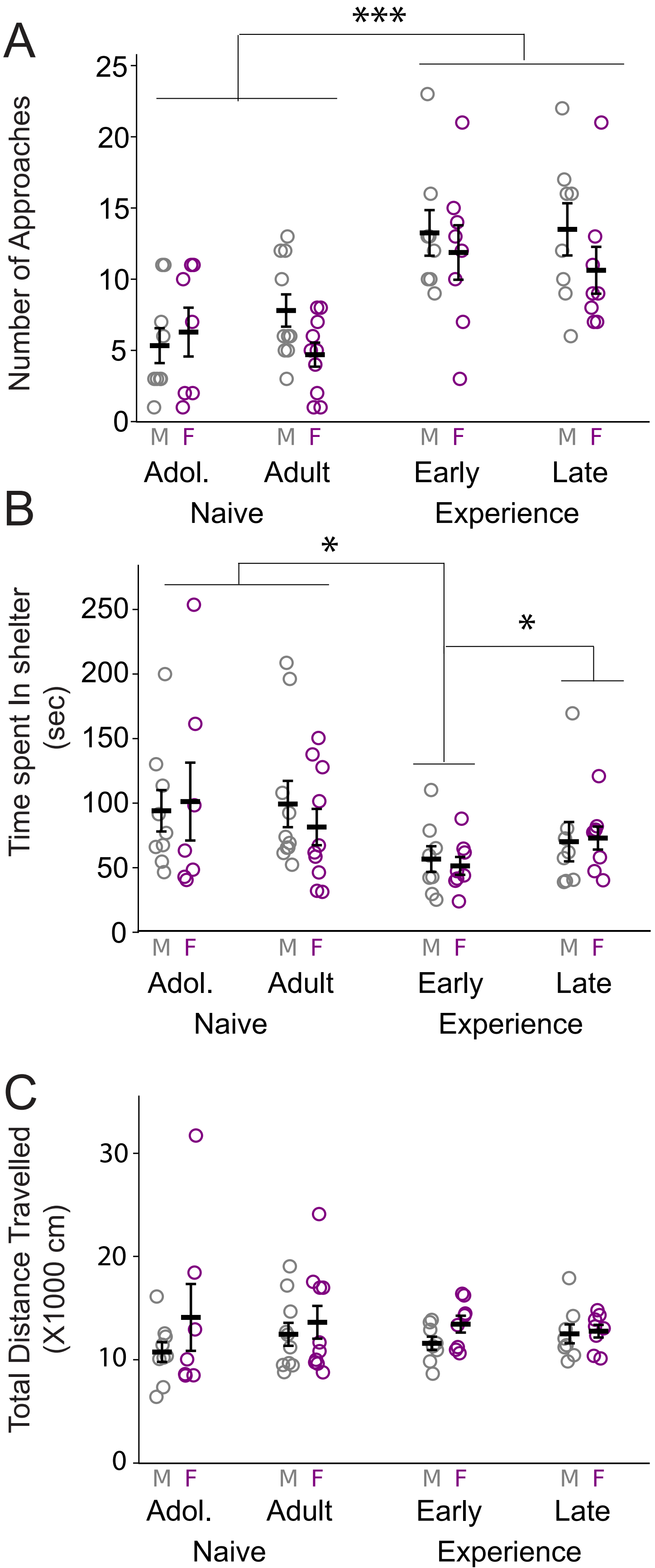

### SUPPLEMENTAL FIGURE 3

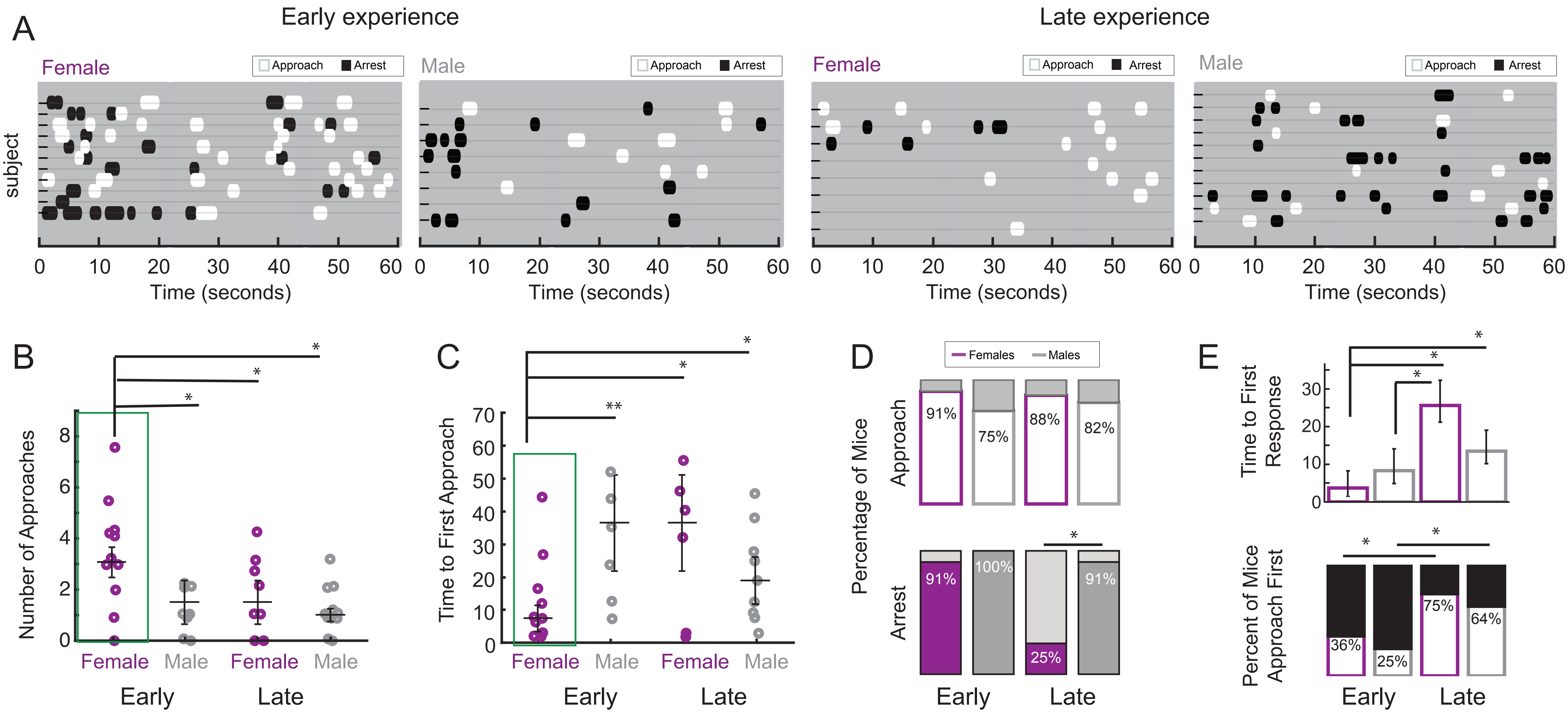

### SUPPLEMENTAL FIGURE 4

A

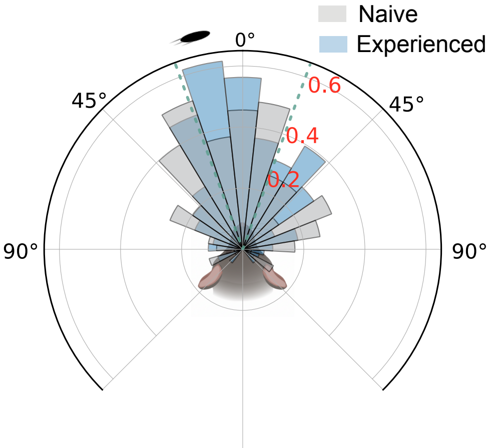

### SUPPLEMENTAL FIGURE 5

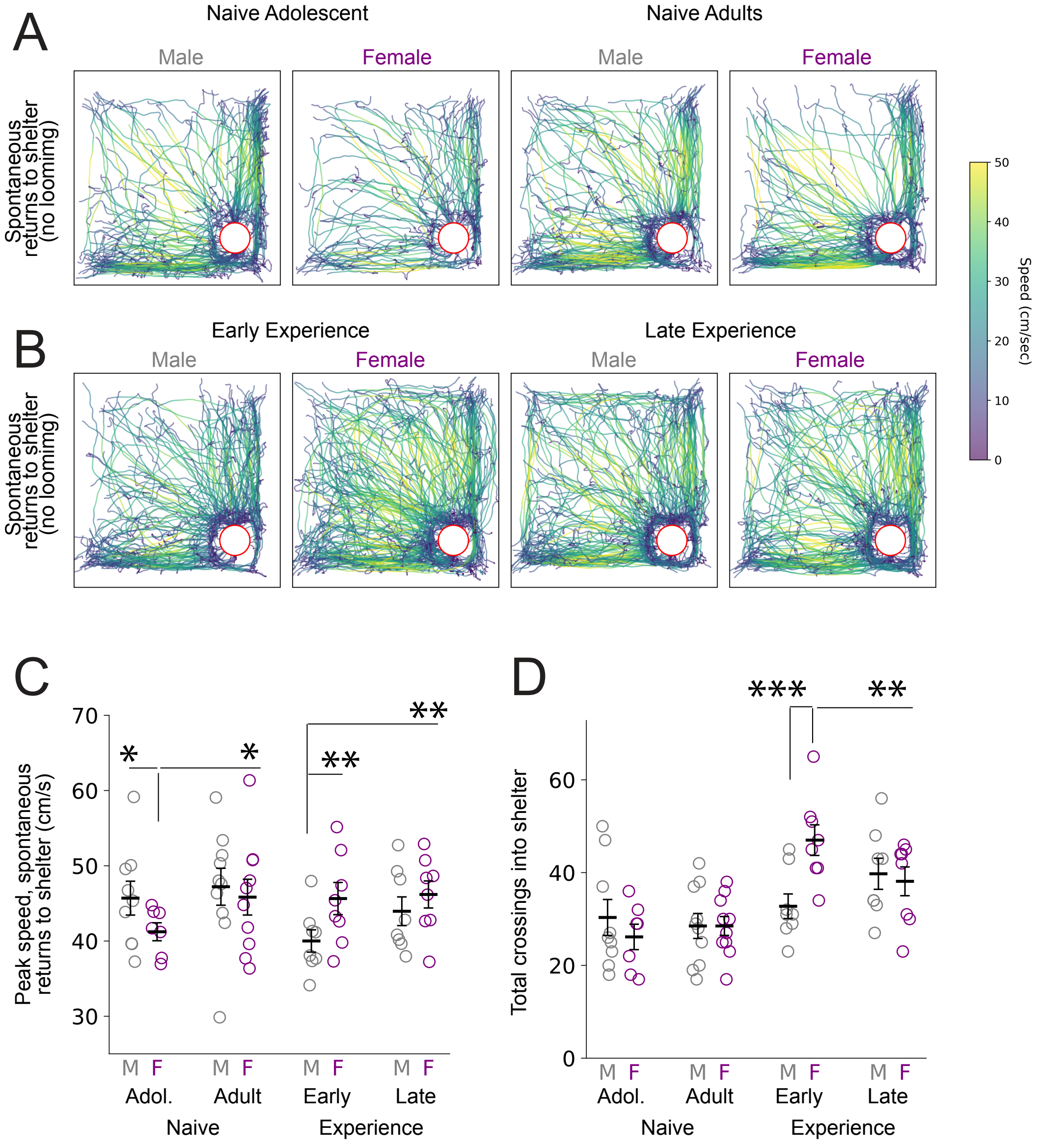
