## Supplemental Figure Legends for "GO, NO-GO: A SENSITIVE PERIOD FOR DEVELOPING SEX-SPECIFIC VISUALLY-GUIDED PREY PURSUIT-PREDATOR AVOIDANCE TRADE-OFF STRATEGIES"

**Supplemental Figure 1: Overview of raw behavior responses to conflicting stimuli by sweeping stimulus valence and trial number.** (A) Ethograms depicting behavior response to the competing valence visual stimulus paradigm for mice without hunting experience tested as adolescents, top, or adults, bottom. The y-axis represents mouse ID, and x-axis represents time in each ethogram. Each 5-minute period is depicted by changes from white to gray shading and then from gray-shading to white, where a different visual target speed (2 cm/sec, 15 cm/sec vs. 50 cm/sec) is presented in each color block. Green, blue and white square markers represent escape, freeze and neither escape nor freeze ‘other’ prey-adjacent response, respectively. Each observable response type is dependent upon the mouse first approaching one of three speeds of sweeping stimulus. (B) Mean escape probability normalized per mouse as a function of number of loom stimulus exposures to determine degree of loom habituation. * = p < 0.05, linear mixed-effects model, N = 9 v. 7 male adolescents versus female adolescents with escapes. No significant differences were found between adult male and females in mean escape probability over trials with loom evoked responses, linear mixed-effects model, N = 10 and 10, adult males and females, respectively. Gray solid lines represent males, and purple solid lines represent females, error bars are standard deviation.  (C) Ethograms depicting behavior response to the competing valence visual stimulus paradigm for mice with hunting experience at different ages, hunting experience as adolescents (early) or as adults (late). Organization and labelling as in A. (D) Mean escape probability as a function of trial number for developmentally timed hunting-experienced mice. Organization and labelling as in B. * = p < 0.05, linear mixed-effects model, N = 8 v. 8 early experience male adolescents versus early experience female adolescents with escapes. No significant differences were found between adult experience male and females in mean escape probability over trials with loom evoked responses, linear mixed-effects model, N = 8 and 8, late experience males and late experience females, respectively.

**Supplemental Figure 2: Basic measures of exploratory and other behavior not immediately associated with loom presentation in our assay.** (A) Mean number of approaches per animal. Experience, regardless of sex or age, enhances sweeping stimulus approach numbers, *** = p < 0.001, main effect of experience, unbalanced, Three-way ANOVA (sex, age, and experience), Tukey’s HSD posthoc testing, N = 9, 7, 10 & 10, naïve male and female adolescent mice, naïve male and female adult mice, respectively, N=8 for all experienced groups. Gray are males, purple females, Error bars are SEM. (B) Mean Time spent in Shelter. Adolescent (early) experience, regardless of sex, leads to decrease in time spent in shelter. * = p < 0.05, interaction between age and experience, Three-way ANOVA (sex, age and experience), Tukey’s posthoc testing, N = 9, 7, 10 & 10, naïve male and female adolescent mice, naïve male and female adult mice, respectively, N=8 for all experienced groups. Gray are males, purple females, Error bars are SEM. (C) Mean total distance travelled during habituation periods. No differences found between groups dependant upon sex, age, nor experience. Three-way ANOVA (sex, age and experience), Tukey’s posthoc testing, N = 9, 7, 10 & 10, naïve male and female adolescent mice, naïve male and female adult mice, respectively, N=8 for all experienced groups. Gray are males, purple females, Error bars are SEM.

**Supplemental Figure 3:  Changes in innate visual orienting responses as a result of developmental stage-specific hunting experience without any loom exposure.** *(*A) Ethograms showing the response of all four groups of mice to sweeping visual motion stimuli in the lower environment. Each row of the ethogram is the response of an individual animal, white indicates time when animals are approaching stimulus, black is when they arrest in response to the stimulus. Each Sweep is outlined by a grey box denoting the time when stimulus is on screen. (B) Mean number of approaches. (C) Time to First approach the stimulus. (D) Percentage of mice in each group exhibiting at least one approach towards sweeping stimulus, white, and those that exhibit at least one arrest in response to the stimulus, black. (E) Mean time to first response indicating their sensitivity to the stimulus at top, and likelihood that they approach the stimulus before arresting (bottom). Significance tested for using a Two-way mixed ANOVA, unbalanced design, sex by age, with Tukey’s HSD post hoc testing for identifying significant differences and to correct for multiple comparison. N =11, 8, 8 & 9 mice, early experienced female, early experienced male, late experienced female and late experienced male groups, respectively. Normality of each dataset was assessed using the Shapiro-Wilk test, Means shown with overall distribution, Error bars are +/- standard error of the mean.

**Supplemental Figure 4: Changes in virtual sweeping stimulus targeting with and without prey capture experience.** Polar plot histogram representing azimuthal angle of approach to visual sweeping stimulus at the onset of overhead looming stimulus for both naive (gray) and hunting experienced (cyan) animals. Visual stimulus angle is significantly different after prey hunting experience. Mean = 18.12 degrees, SEM = 3.89 degrees versus mean = 38.20 degrees, SEM = 4.77 degrees, p < 0.01, Watson’s U, N = 205 and 383 approaches for naïve versus experienced animals, respectively.

**Supplemental Figure 5: Measures of darting behavior between shelter and other areas of the environment not immediately associated with loom presentation.** (A) Trajectories of Naïve adolescent versus naïve adult mice returning to shelter outside of the immediate loom presentation period (> 5 seconds post loom or before first loom). Each trajectory is color coded by mouse locomotion speed, color bar shown to the right. The location of the shelter is indicated by red open circle in lower right corner of each plot. (B) Trajectories derived from spontaneous returns to the shelter for prey capture experienced mice that hunted at different developmental stages, adolescent (early) versus adult (late). Color coding and representations as in A. (C) Mean peak speed of the spontaneous returns to shelter. Running speeds to shelters apart from looms is dependent on an interaction between sex, age and age of experience. * = p < 0.05, ** = p < 0.01, unbalanced, Three-way ANOVA, N = 9, 7, 10, 10, 8, 8, 8, 8, naive adolescent male, naive adolescent female, naive adult male, naive adult female, early experience male, early experience female, late experience male, late experience female, respectively. Error bars are SEM. (D) Mean total crossing into shelter for all groups. Escaping to shelter was dependent on an interaction between sex, and age of experience. ** = p < 0.01, *** = p < 0.001, unbalanced, Three-way ANOVA, N = 9, 7, 10, 10, 8, 8, 8, 8, naive adolescent male, naive adolescent female, naive adult male, naive adult female, early experience male, early experience female, late experience male, late experience female, respectively. Error bars are SEM.
